# ro-crate-rs: Development of a Lightweight RO-Crate Rust Library for Automated Synthetic Biology

**DOI:** 10.64898/2026.01.22.701040

**Authors:** Matt Burridge, Zhen Ou, Katherine James, Gizem Buldum, Jesmine Lim, James Finnigan, Simon Charnock, Anil Wipat

## Abstract

Advances in laboratory automation and AI-driven experimental design have increased the scale and complexity of data generated in synthetic biology. Whilst biofoundries provide significant resources and infrastructure to execute these experiments, most laboratories rely on isolated automated instruments and software systems that operate as disconnected silos, producing heterogeneous data formats with little structured metadata. This fragmentation hinders data integration, reproducibility, and downstream computational workflows.

A potential solution is RO-Crate, which offers a lightweight, extensible framework for packaging research data with machine-readable metadata, but existing tooling remains immature for automation-orientated, cloud-native, or high-throughput laboratory workflows. Here, we introduce **ro-crate-rs**, a new suite of tools centred on a performant Rust library for constructing, validating and packaging RO-Crates across diverse compute environments and automated hardware. The library enforces RO-Crate 1.1 constraints through strong typing while enabling flexible extensions, and is complemented by a Python API and CLI for interactive use and pipeline integration.

We demonstrate this combined approach through a semi-automated Old Yellow Enzyme characterisation workflow, showing how RO-Crates can capture data and metadata across multiple independent instruments. Together, these tools provide a robust foundation for FAIR-compliant, automation-ready data management and enable reproducible reconstruction of experimental workflows even in non-biofoundry settings.

**Availability:** https://github.com/intbio-ncl/ro-crate-rs

## Introduction

Automation has become increasingly essential in synthetic biology as the scale and complexity of experimental workflows continue to grow.^1^ Although automation is often associated with liquid-handling systems, modern laboratories rely on a broad range of automated instruments, including microscopes, plate readers, incubators, colony pickers, and thermocyclers. These devices can be integrated into workcells and, in some cases, combined into fully automated facilities known as biofoundries.^2^ Biofoundries aim to minimise human involvement in the experimental design-build-test-learn (DBTL) cycle by linking laboratory hardware to computational platforms, data-management systems, and workflow schedulers.^1,3–5^

Despite their advantages, biofoundries remain relatively uncommon; they require substantial capital investment, stable funding, specialised infrastructure and expertise, and long-term maintenance.^6^ Consequently, most laboratories operate isolated workcells or standalone automated instruments rather than fully integrated facilities. This fragmentation creates two major challenges. First, automated systems are difficult to program, maintain, and adapt without specialised expertise. Second, the data they generate are often stored in proprietary formats or vendor-specific log files, making extraction, integration, and reuse cumbersome. One can think of these problems as split between automated experiment execution and automated data acquisition, both of which are essential challenges for reproducible synthetic biology.^7^ Fundamentally, a sufficiently described experiment that is implemented on an automated platform should be able to be reconstructed from its data and metadata, supplanting the need for describing what should have happened, by precisely recording what actually happened.

Efforts are emerging to address these issues. For protocol execution, the growing lab-automation community provides guidance and open-source tools, most notably *pylabrobot*, which aims to offer a unified Python API for interacting with diverse automation hardware.^8^ Additionally, for machine-readable protocol description, *LabOP* is in development^9^ and for hardware integration, SiLA 2 is emerging as the dominant standard.^10^ In parallel, tools such as *Parsley*^11^ and *Flapjack*^12^ support parsing and analysing plate-reader outputs, and recently there has been collaboration within the Global Biofoundry Alliance^13^ to standardise workflows within biofoundries.^14^ However, to our knowledge, there is still no general solution for structuring data produced by siloed automated hardware in a way that facilitates downstream dry-lab workflows.

## Background

RO-Crate offers a promising foundation for addressing this gap. RO-Crate is a lightweight metadata description framework that enables a researcher to package their data with the required amount of structured, machine-readable metadata.^15^ It is based on JSON-LD^16^, a linked-data format, allowing both local and remote files to be described in a machine-readable format that is backed by the minimal vocabulary of schema.org.^17^ Additional vocabularies can then be included if the research data requires it, for example using Bioschemas for biological metadata^18^, or if the RO-Crate is being used to describe in a highly domain-specific use case, such as a Nextflow workflow^19^, it can conform to a defined **RO-Crate profile**.^20^ The goal is to provide a comprehensive yet accessible description framework that enables researchers to conform to FAIR data principles^21^ with minimal overhead.

One of the key strengths of RO-Crate is its core simplicity and extensibility, allowing it to be used throughout many research domains. This flexibility, however, creates a need for domain-specific tooling to support specialised use cases, including those in synthetic biology where interoperability and reproducibility remain ongoing challenges. RO-Crate can describe data generated across cloud environments, local computing infrastructure, or proprietary lab instruments, offering a way to bridge disconnected data sources without requiring heavyweight data-management platforms.

In this work, we present a new suite of RO-Crate tools designed to meet the needs of computational and experimental synthetic biologists. At its core is a lightweight, performant Rust library for creating, modifying, and validating RO-Crates, leveraging the Rust type system and the ‘serde’ ecosystem to enforce RO-Crate 1.1 constraints with strong data handling and reliability. On top of this core, we provide a Python API for convenient, ad hoc crate generation and a command-line interface for basic crate manipulation and validation. Together, these tools enable RO-Crate integration across cloud platforms, lab workstations, and automated instrumentation.

## Results

### Core ro-crate-rs Library

The top level of the RO-Crate JSON-LD structure is a **@context** and a **@graph** object. Within **@context**, a reference to another JSON-LD object which contains RO-Crates own schema.org vocabulary is defined (allowing RO-Crate specific versioning), as well as any other domain-level contexts the RO-Crate requires. The **@graph** object is an array of other objects, with each object conforming to a particular **entity** as described in Table 1.

**Table 1:** Entity types found within an RO-Crate.

| Entity Type | Occurrence | Description |
| --- | --- | --- |
| Metadata Descriptor | One | A self-describing object that defines the ro-crate-metadata.json file as a CreativeWork that conforms to the RO-Crate specification. It also defines the Root Data Entity. |
| Root Data Entity | One | The root directory object (i.e "./") that contains the RO-Crate Metadata File (ro-crate-metadata.json), which describes the data within the crate |
| Data Entity | Many | An object that describes a file or a directory stored locally or remotely |
| Contextual Entity | Many | An object that defines metadata that is used to describe data entities |

The ro-crate-rs library defines this structure within a hierarchical data structure of type ‘rocrate’ which contains a ‘context’ of type ‘RoCrateContext’ and ‘graph’ which is a vector of type ‘GraphVector’.

Within ‘RoCrateContext’ you can define one of the three following types; ‘reference’, ‘embedded’, ‘extended’.

- A reference context is generally considered to be the default of an RO-Crate, with a String of https://w3id.org/ro/crate/1.1/context.
- An embedded context is a hash map consisting of key:values describing vocabulary used. This is rarely used as if a context conforms and is deserialised to this type, the crate would not conform to v1.1 specification.
- An extended context combines both the reference context, defining the RO-Crate v1.1 context, as well as a vector of key:value pairs that describe more domain specific properties that have been used within the crate.

Within ‘GraphVector’ you can define one of the following; ‘MetadataDescriptor’, ‘RootDataEntity’, ‘DataEntity’, ‘ContextualEntity’ (Figure 1).

**Figure 1:**
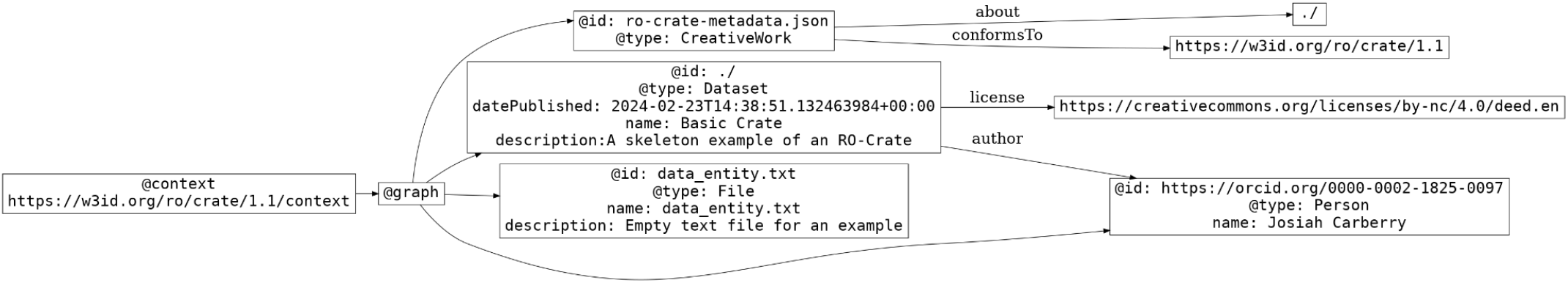
Visualisation of an RO-Crate with basic data. The **@context** (’RoCrateContext’) signifies that the RO-Crate uses the v1.1 vocabulary. Within the **@graph** (’GraphVector’) from top to bottom, there is a **Metadata Descriptor** (’MetadataDescriptor’), **Root Entity** (’RootDataEntity’), **Data Entity** (’DataEntity’) and a **Contextual Entity** (’ContextualEntity’). The metadata descriptor defines its root directory as an RO-Crate (’about’), pointing towards the v1.1 specification context (’conforms_to’). The root entity describes the overall crate, giving it a name, description, and stating when it was published. Within the root entity, the data license (’license’) for the overall is also defined. The data entity describes the ‘data_entity.txt’ file. The contextual entity describes a person, who is also the author of the RO-Crate.

- A metadata descriptor defines the root of the RO-Crate as ‘./’, as well as confirming that the crate uses the RO-Crate v1.1 context.
- A RootDataEntity describes the crate and the root ‘./’. This is where the overarching reason of the crate is described, as well as linking all files that need to be described within the crate.
- A ‘DataEntity’ describes a data entity within a crate, a metadata object that describes data which is contained within the crate. For example, this could be a CSV file, image or FASTA file.
- A ‘ContextualEntity’ describes a contextual entity, a metadata object that defines metadata surrounding either the crate itself, or metadata for describing a ‘DataEntity’ with more context.

Both the ‘DataEntity’ and ‘ContextualEntity’ currently function the same, and they are more conceptually defined. A ‘DataEntity’ is an entity of data which has been produced and/or used in the research that the crate describes. The entity itself needs to point to a valid file path for it to be deserialised into a data entity. A ‘ContextualEntity’ is an entity which helps describe a ‘DataEntity’, ‘RootDataEntity’ or ‘MetadataDescriptor’. It can be said that as itself, the data it may contain is not required to understand the crate, but more helps provide additional clarity or context to particular entities within the crate. These ‘ContextualEntities’ may be valid URLs, and they may lead to datasets or files that are stored remotely.

Each entity structure consists of an ‘@id’ (or ‘id’ for struct field) and a ‘@type’ (or ‘type_’ for struct field). For ‘MetadataDescriptor’ and ‘Root Data Entity’, they consist of other fields that have been determined to be a requirement by the RO-Crate specification. Fields that have been determined as **MUST** by the RO-Crate specification are required in their relevant data structure, whilst fields determined as **SHOULD** are Options. Each entity structure also has a field called ‘dynamic_entity’, which allows the population of any form of JSON-LD compatible value within the statically typed requirements of Rust. These, by default, are instantiated as ‘None’. Any field that is not described as **MUST** or **SHOULD** within the specification can be added as a ‘dynamic_entity’.

A design choice of this library is to avoid introducing abstractions beyond the RO-Crate specification. Rather than providing high-level schema.org classes (e.g., ‘Person’, ‘Dataset’), the library exposes only the four RO-Crate entity types. This keeps the implementation transparent and avoids enforcing domain semantics.

### Reading and Writing

Reading and writing are based on Serde, a serialisation/deserialisation framework. All core RO-Crate structs implement customised Serde JSON traits, enabling direct conversion between valid RO-Crate JSON-LD and the internal ‘RoCrate’ data structure.

The library supports reading crates from:

- any RO-Crate 1.1–compliant JSON file
- a raw JSON string
- a zipped archive containing a ‘ro-crate-metadata.json’ file

This allows large packaged research objects to be parsed without full decompression, reducing unnecessary I/O.

### Validation

Validation is currently implemented in two ways. First, the strongly typed data structures enforce the RO-Crate 1.1 specification by requiring all **MUST** properties and restricting values to valid JSON-LD datatypes. This prevents malformed structures, unexpected nesting, or illegal values.

The second way is during crate reading, where three validation levels can be toggled depending on context. The first level allows crates to be read with base validation, where if the crate is in the minimal structural form of an RO-Crate it will successfully load. The second level enables warnings, whereby if the crate uses types and properties that are not defined within the schema.org vocabulary, these are flagged. The third level is strict, where if the RO-Crate does not conform in structure nor vocabulary, it is not loaded. This approach reduces the chance of non-compliant crates propagating through workflows while keeping flexibility for exploratory use.

### Packaging

RO-Crates are intended to function as portable, self-contained research objects, so packaging support is built into the core library. The packaging process preserves data-entity identifiers and automatically rewrites file paths as relative paths within the archive. Users may retain the directory structure, which is useful for profile-specific layouts such as the ISA data model^22^, or flatten it for downstream processing. Additionally, data entities that are referenced that are external to the Root metadata directory can be imported for an entirely self-contained package.

To ensure portability, packaging automatically adds a ‘@base’ UUID URN to the crate’s ‘@context’, providing stable, valid URIs for all referenced entities even after extraction or relocation.

### File Conversion

Whilst leveraging the JSON format enables complete flexibility to what is described, there are situations where alternative, tabular formats are required. Currently, this library allows easy conversion to two tabular formats, CSV and Apache Parquet. Tabular conversion follows a quasi N-Quads structure^23^, with Graph (G), Subject (S), Predicate (P) and Object columns, allowing for more efficient storage within archival data stores.

### Python API

While the Rust library provides a performant and strongly typed core, many users need to construct RO-Crates in interactive or ad hoc environments. Python remains the dominant language in scientific computing, particularly within the synthetic biology community, so the Python API is designed to provide convenient access to the same robustness as the Rust implementation.

The API mirrors the Rust entity model but uses plain Python dictionaries and a small set of simple functions for crate construction. Users can assemble entities as basic dicts while relying on the underlying Rust core for structural validation, context handling, and conformance to the RO-Crate 1.1 specification.

Unlike the higher-level ‘ro-crate-py’^24^ and pydantic-based^25^ libraries, this API does not guide users through schema-specific classes. Instead, it offers a flexible interface that assumes some familiarity with the RO-Crate structure. This design intentionally balances flexibility with strictness: users may choose their own level of semantic detail, while the core library enforces the required structural constraints.

### CLI tool

The CLI tool provides lightweight, script-friendly access to core functionality, enabling users to create, inspect, modify, validate, and save RO-Crates. It is not intended for rapid or large-scale crate construction, but rather for interacting with existing crates in constrained environments such as headless servers, automated pipelines, or shell scripts. Typical uses include structural validation, quick inspection, and basic type checking against schema.org terms.

### Enzyme Characterisation Use Case

To demonstrate how the toolkit can structure data across heterogeneous laboratory processes, we outline a simplified enzyme-activity characterisation workflow. In this example, a panel of ene-reductases (Old Yellow Enzymes; OYEs) are purified, undergo a FRED assay^26^, and are quantified on a plate reader for catalytic activity analysis (Figure 2).

**Figure 2:**
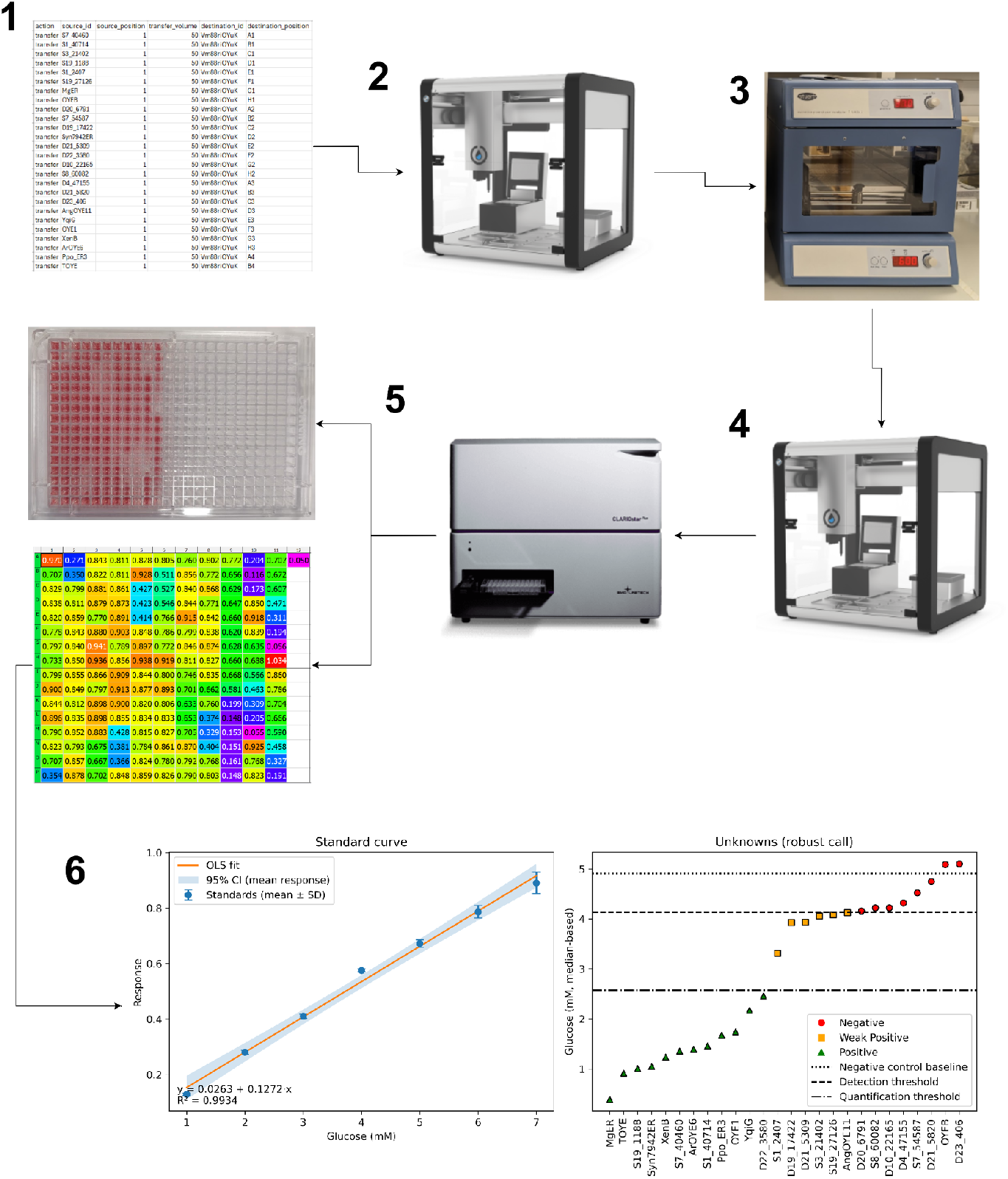
Example use case for a minimalistic RO-Crate based data capture for automated experimental workflows. (1) Denotes 96 well plate microplate setup of manually purified Old Yellow Enzymes. (2) Enzyme plate is used for a biotransformation setup to test catalytic activity on target substrates. (3) Overnight incubation. (4) GOD-POD Assay setup to measure glucose concentrations. (5) Plate reader absorbance 510 nm analysis. (6) Analysis of raw plate reader to determine statistically relevant positive/negative enzyme activities. All automated liquid handling was executed on the Opentrons OT-2. Plate reader analysis was carried out on BMG Clariostar.

Each major experimental step produces its own RO-Crate, all of which have been packaged and archived in Zenodo (https://doi.org/10.5281/zenodo.17828765), resulting in six crates that collectively describe the full workflow:

1. Microplate setup
2. OT-2–executed biotransformation
3. Overnight incubation
4. OT-2–executed GOD-POD colorimetric assay
5. BMG Clariostar plate-reader data acquisition
6. Statistical analysis of enzyme activity

RO-Crate 1 describes the initial wet-lab setup. It includes a CSV mapping file identifying the well locations and enzyme variants, along with a protocol detailing transfer of purified enzymes to target plate and aliquoting into a 96-well plate.

RO-Crate 2 captures the automated biotransformation step performed on an OT-2 liquid-handling robot, documenting how enzyme aliquots were transferred into the reaction mixture.

RO-Crate 3 represents the incubation phase, providing metadata about the shaking incubator conditions and linking the input and output plates.

RO-Crate 4 records the GOD-POD colorimetric assay performed on the OT-2, detailing reagent dispensing and plate handling to measure OYE activity.

RO-Crate 5 contains the absorbance data generated on a BMG Clariostar plate reader, enabling quantitative analysis of enzyme activity for each well.

RO-Crate 6 contains a notebook which analyses the raw data generated from the plate reader analysis in RO-Crate 5, and runs statistical tests to determine whether the tested enzymes have statistically significant catalytic activity.

Together, these six crates provide a modular, machine-readable description of the entire workflow, illustrating how RO-Crates can integrate data across disconnected instruments and automation steps without the need for a substantial data infrastructure. Additionally, these crates provide essential data and context for LLM interpretation for future agentic workflow analysis, as well as the reconstruction of exact protocols for experimental reproducibility.

## Conclusion and Future Direction

The ro-crate-rs library provides a robust core for further developing ro-crate tooling and enforces the RO-Crate specification at read, modification and write. The Rust library enables performant, process-orientated RO-Crate generation, while the Python library enables ad hoc modification and RO-Crate utilisation in standard data pipelines with a very simple API. Similarly the CLI tool can be used for ad hoc RO-Crate utilisation as well as being implemented in Bash workflows.

Whilst this technical note describes the initial release of the library, there is significant progress to be made. The next release will include RO-Crate Profile handling and further user testing is required to ensure it is a suitable fit for widespread use. Currently it is designed around the v1.1 specification, and with the recent release of v1.2 into long-term support, updates will have to be made. Implementation of external context validation is required, and investigations into whether further built-in validation is worthwhile need to be carried out. Additionally, direct object store streaming needs to be implemented to allow cloud-native RO-Crate reading.

For the automation context, more work needs to be carried out on stateless data handling and interoperability of basic lab data models. For instance, there is no accepted ontology nor data model for basic labware, such as microplates or microcentrifuge tubes. There is also no defined data model for describing liquid handling executions, with no defined semantics for aspirations, dispenses, tip pickups or ejections. As such, work needs to be carried out in these fundamental areas that will enable true FAIR data within automation enabled labs.

## Data Availability

ro-crate-rs is a fully open source library that can be accessed at https://github.com/intbio-ncl/ro-crate-rs. All data generated in the use case can be found in Zenodo (https://doi.org/10.5281/zenodo.17828765). The Rust library is available in crates.io (http://crates.io), along with the CLI tool (https://crates.io/crates/ro-crate-rs and https://crates.io/crates/ro-crate-rs-cli). Additionally, the Python library is available at PyPI (https://pypi.org/project/rocraters/).

## Acknowledgements

The authors would like to thank Dr. Soiland-Reyes, Mr Owen and Mr Bacall for their initial insights into using RO-Crate for lab automation data handling. Funded by BBSRC Bioscience for Sustainable Consumer Products CTP.

